# Port of Protein-Protein Interactomes: An experiment-based protein-protein interactome database for rice

**DOI:** 10.64898/2026.08.16.744343

**Authors:** Xixi Liu, Jiawen Lu, Lili Jia, Dandan Xia, Jie Huang, Yu Cheng, Mengyuan Li, Yiting Chen, Xinyong Liu, Guanghao Li, Wanning Liu, Jia Li, Jiezheng Ying, Yifeng Wang, Zhiyong Li, Xiaohong Tong, Yuxuan Hou, E Zhiguo, Jianwei Zhang, Jian Zhang

**Affiliations:** State Key Lab of Rice Biology and Breeding, China National Rice Research Institute, Hangzhou 311400, China; National Key Laboratory of Crop Genetic Improvement, Hubei Hongshan Laboratory, Huazhong Agricultural University, Wuhan 430070, China

**Author notes:** Corresponding author: Prof. Jian Zhang; *E-mail address:*; Prof. Jianwei Zhang; *E-mail address:*; Associate Prof. Zhiguo E; *E-mail address:*. These authors contributed equally (Xixi Liu, Jiawen Lu, Lili Jia, Dandan Xia, Jie Huang).

## Abstract

Protein–protein interactions (PPIs) play a crucial role in enabling proteins to carry out their functions within various biological processes (Hui et al., 2003). Since the introduction of the yeast two-hybrid (Y2H) method for PPI detection in 1989 (Fields and Song, 1989), the identification of PPIs has become a significant focus in modern biological research. PPI goes beyond examining individual proteins, allowing researchers to establish a comprehensive network that regulates biological processes. Rice, as a key model organism in plant biological studies, has been at the forefront of PPI research. In 2008, prominent rice scientists in China called for concerted efforts to define a comprehensive protein–protein interaction network experimentally, which aimed to facilitate the prediction of the functional mechanisms operating throughout a plant’s lifecycle (Zhang et al., 2008). With efforts for 2 decades, the experimentally identified rice PPIs have reached over ten thousand. Several public databases have been established to systematically collate and store PPIs, including STRING (Szklarczyk et al., 2019), BioGRID (Oughtred et al., 2020), IntAct (del Toro et al., 2022), PRIN (Gu et al., 2011), RicePPINet (Liu et al., 2017) and RiceNet v2 (Lee et al., 2015). However, most PPI datasets in rice stem from computational predictions, while experiment-based rice PPI datasets are fragmented due to the lack of systematic profiling at the rice PPIome level, which largely hinders information sharing in the rice research community.

To bridge this gap, we constructed the Port of Protein-Protein Interactomes (POPPIN; https://riceome.hzau.edu.cn/poppin/), an integrated database dedicated to sharing experimentally verified PPIs and functional clues in rice. Empowered by high-throughput PPIome profiling technologies and text mining assisted by a large language model (Huang et al., 2025; Liu et al., 2025), POPPIN currently has deposited over 150,451 pieces of rice PPI-related information. Additionally, POPPIN provides detailed protein information, including GO annotations, subcellular localizations, domains, trait ontology (TO) information, and hyperlinks to external biological databases. Through offering a user-friendly web interface for search and dynamic network visualization, POPPIN serves as the first large-scale, experiment-based database for searchable PPIs in rice, and has the potential to be extended to other species under this structural framework.

## PPI Data sources

Currently, POPPIN has deposited two rice-related PPI datasets, namely PPIs within rice proteins (rice × rice) and PPIs between rice and *Arabidopsis thaliana* proteins (rice × Arabidopsis). Based on the data sources, the rice × rice PPI dataset could be divided into five catalogs (Figure 1A; Supplemental Table 1). (i) 81,604 randomly identified PPIs by the yeast two-hybrid (Y2H) system. Through mating the Y2H bait cDNA library and prey cDNA library, random positive PPI colonies were selected on SD/-Leu-Trp-His-Ade/+5 mM 3-AT medium and further profiled by BIP-seq or TDOP-seq technologies (Figure 1A; Supplemental Table 1) (Huang et al., 2025; Liu et al., 2025). (ii) 42,444 randomly identified PPIs by bacterial two-hybrid (B2H), which follows the same pipeline as (i), but in an E.*coli* system under the selection of SD/+M9/-His/+5 mM 3-AT/+Stre/+Chl/medium (Figure 1A; Supplemental Table 1). (iii) 11,977 PPIs extracted from publications by text-mining. To this end, 23,000 full-text articles and 27,650 abstracts were fed into a large language model (LLM)-assisted literature-mining workflow (Figure 1D-F), where the texts were chunked into manageable blocks, and DeepSeek-V4-Flash was optimized using a benchmark set of 100 articles through systematic prompt and parameter tuning. At a temperature of 0.1, the workflow achieved a precision of 86.76% (354/408), a recall of 80.63% (354/439), and a calculated F1 score of 83.59%. (iv) 2,764 confirmed non-PPI pairs. Through pairwise mating of manually constructed 37 baits and 147 preys, we identified PPIs for 5,439 combinations, which finally yielded the 2,764 non-PPI pairs (Figure 1A; Supplemental Table 1). (v) 4,964 auto-activator proteins in the Y2H system. By either mating a bait cDNA library with an empty pGBKT7 vector, or a prey cDNA library with an empty pGADT7 vector, proteins with auto-activities in the Y2H system could be selected and identified by pac-bio sequencing (Figure 1A; Supplemental Table 1). For the rice × Arabidopsis dataset, 7,726 PPIs covering 1,270 rice proteins and 2,144 Arabidopsis proteins were identified by the Y2H system coupled with TDOP-seq technology (Figure 1A; Supplemental Table 1) (Huang et al., 2025). In total, 150,451 pieces of rice PPI-related information covering 1,270 rice proteins and 2,144 Arabidopsis proteins are provided in POPPIN.

**Figure 1.**
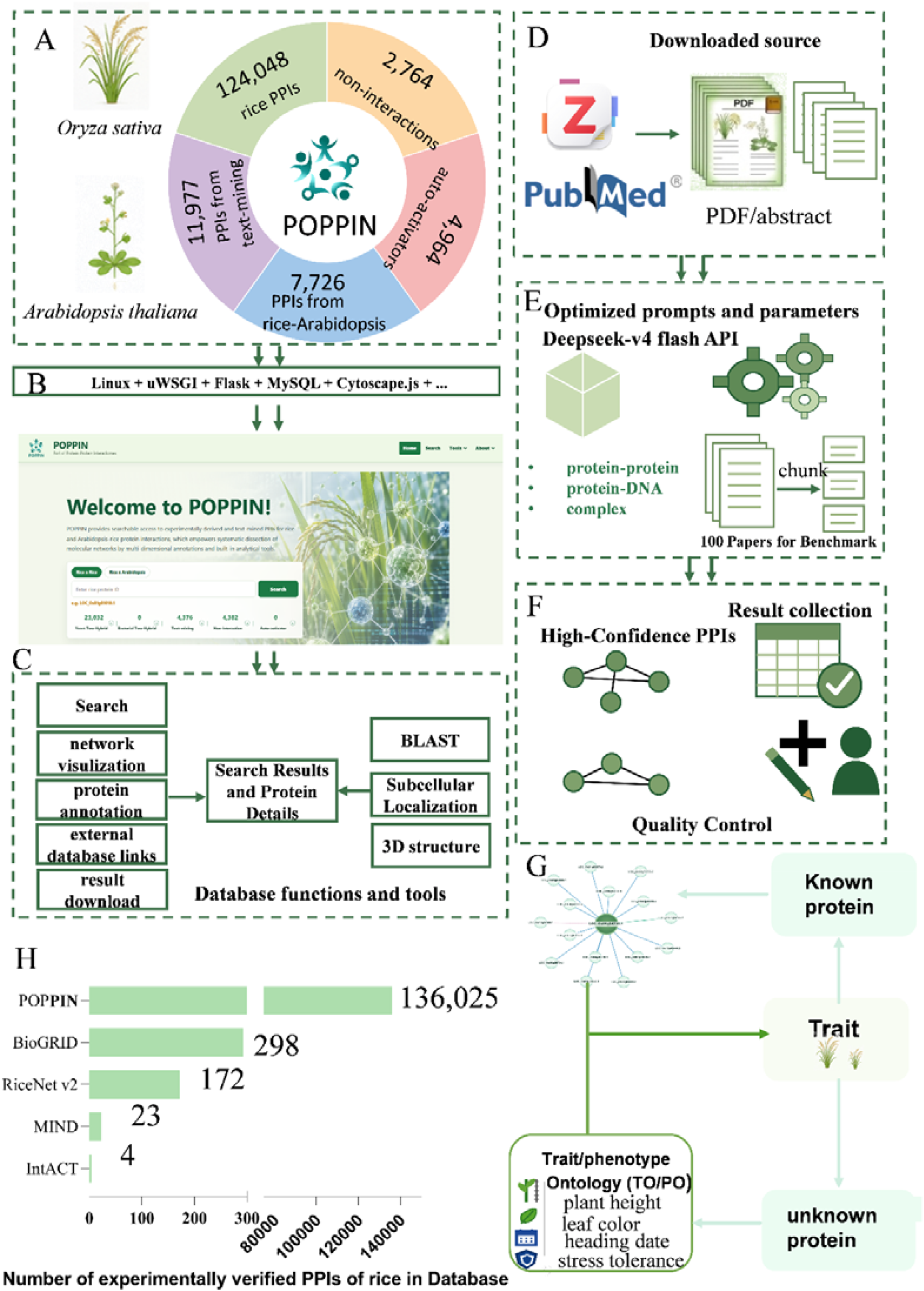
POPPIN database **(A-C)** Overview of the construction of POPPIN Database. (A) The summary of the POPPIN database content and evidence categories. (B) The layer shows the tools used to build the POPPIN website. (C) The POPPIN architecture shows the functions and tools. (D-F) The workflow of text mining. (D) Relevant publications were collected manually or through Zotero or PubMed. (E) Large language model-assisted interaction extraction. (F) Result collection and verification. (G) The example of biological application of POPPIN. (H) Comparison of experimentally verified rice PPIs among POPPIN and other reported databases.

### Structure of the database

POPPIN is a comprehensive platform designed to systematically analyze and visualize PPIs in rice, consisting of a data layer, an application layer, and a presentation layer (Figure 1B-C). POPPIN is hosted on a Linux operation system and a Nginx web server. The data layer stores standardized PPIs, interaction records, evidence channels, and functional annotations in a MySQL database. On the client side, users query and download interaction records via a Python/Flask-based backend. The presentation layer provides search interfaces, interaction tables, network visualization, detailed protein and interaction information, and links to external biological resources. ECharts (Deqing et al., 2018) and Cytoscape.js (Max et al., 2015) are used for interaction-network visualization.

### Searching PPIs in POPPIN

POPPIN offers both basic and advanced PPI searches. From the home page, users can search the database within the selected dataset by gene ID or gene name using the basic search box (Figure S1). POPPIN accepts multiple inputs and flexible rice gene IDs and reported gene names. For example, SAPK10, OsSAPK10, LOC_Os03g41460 (MSU ID), and Os03g0610900 (RAP ID) are well compatible for searching. Alternatively, users can perform an advanced search on the “search” webpage, where the job could be customized by selecting experiment types and the interaction hierarchies (Figure S2 A). POPPIN supports up to 3 hierarchies, in which a PPI network of “Protein A->Protein B->Protein C->Protein D” could be presented for the input protein A (Figure S2 A).

On the results page, the PPI network for the input protein(s) is graphically visualized by nodes and edges. Variations in edge colors and styles indicate different experiment types. Users can customize node labels to display gene IDs, gene names, or both in the network graph. Additionally, the graph can be downloaded in PNG format (Figure S2 B). Clicking on the nodes or edges reveals further details about the objects, including annotations, subcellular localization, conserved domains, PPI types, and supporting evidence, which are displayed in a window adjacent to the graph (Figure S2 B). A downloadable CSV table containing information about the interactive proteins is also available (Figure S2 B). Trait ontology annotations provide functional insights into the target proteins (Figure 1G; Figure S2 C). To enhance accessibility, both gene names and annotation entries are linked to external databases such as Rice Gene Database Based on INTERNET (Rice Data; https://www.ricedata.cn/gene/index.htm) (E et al., 2006), Rice Gene Index (RGI; https://riceome.hzau.edu.cn/) (Yu et al., 2023), Rice Genome Annotation Project (RGAP; https://rice.uga.edu/) (Hamilton et al., 2025), The Rice Annotation Project Database (RAP-DB; https://rapdb.dna.naro.go.jp/) (Kawahara et al., 2026), National Center for Biotechnology Information (NCBI; https://www.ncbi.nlm.nih.gov/), and Bing (Microsoft Bing; https://cn.bing.com/) (Figure S2 C).

Moreover, POPPIN features user-friendly built-in tools, including BLAST, subcellular localization analysis, and a 3D structure viewer (Figure S3-S4).

In summary, POPPIN is not merely a repository for rice PPI data; it also encompasses positive experimental PPIs, negative interaction tests, auto-activator controls, cross-species PPIs, and serves as a vital connection between interaction maps and phenotypic biology in rice. Although it currently focuses on rice, there is potential for the POPPIN database to expand to include inter-species PPIs, such as those involving rice × pathogens and rice × insects, and ideally, other species beyond rice as well. This expansion is anticipated to enhance functional genomics and facilitate inter-species gene function transfer by linking genotype to phenotype through the physical interactome.

## Supplementary methods

### Text-mining

To complement the experimental datasets, we performed large language model-based mining of scientific literature to get additional interactions. We downloaded abstracts and full-text articles related to rice or *Arabidopsis thaliana* using Zotero combined with manual methods. The NCBI query was the following query= ((“rice” [Title/Abstract] OR “oryza sativa”[Title/Abstract] OR “Arabidopsis” [Title/Abstract]) AND (“interaction”[Text Word] OR “interact” [Text Word] OR “interacted”[Text Word] OR “ubiquitinate”[Text Word] OR “phosphorylate”[Text Word])). In total, 23,000 articles and 27,650 abstracts were downloaded for analysis.

To efficiently get interactions from literature, we systematically optimized the DeepSeek□V4□Flash model. Prompts and parameters were fine-tuned using a benchmark set of 100 articles to optimize the workflow for extracting accurate pairs. The evaluation metrics were calculated as follows:

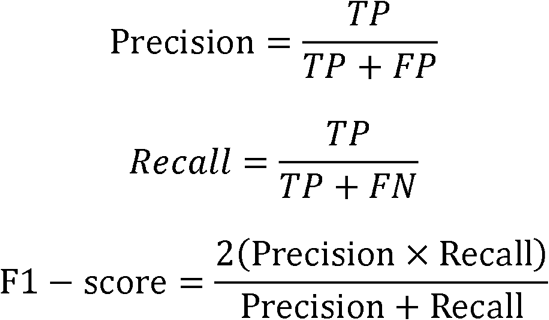

*TP, FP*, and *FN* correspond to true positives, false positives, and false negatives, respectively.

The temperature coefficient was set strictly to 0.1 to suppress model hallucinations and maximize accuracy. PDF texts were chunked into appropriate blocks by dedicated Python scripts. The DeepSeek□V4□Flash API was subsequently executed at scale to extract and summarize interaction data. Extracted records were standardized into a unified format containing Interactor1, Interactor2, Relation, Method, Evidence, and PDF_doi. All candidate interactions were subsequently filtered and manually curated by experts, yielding a high-confidence, literature-derived PPI dataset.

### Architecture and data source

POPPIN is an interactive resource website based on the Linux operating system and the Nginx web service. The current web application is implemented in Python using Flask and Jinja2, and is served through uWSGI or Gunicorn. Protein–protein interaction and gene annotation records are stored in a MySQL database and accessed using PyMySQL and a DBUtils connection pool. Additional annotation indexes, BLAST indexes, caches, and data snapshots are maintained in SQLite and local JSON, CSV, and Excel files. The front end is developed using Bootstrap, jQuery, Popper.js, and DataTables. ECharts and Cytoscape.js are used for interaction-network visualization. BLAST+, Mol*, SwissBioPics, UniProt, and AlphaFold DB support sequence search, protein annotation, localization, and structure visualization.

TO annotations were downloaded from Rice Data and TAIR. The domain data in rice was downloaded from CDD: https://ftp.ncbi.nih.gov/pub/mmdb/cdd/cdd.tar.gz. Cellular localization data were downloaded from the website: https://wolfpsort.hgc.jp.

### POPPIN benchmarking and Validation

To ensure dataset accuracy, POPPIN employs rigorous validation strategies. To validate the PPIs, large-scale Y2H and E2H datasets performed assays. Specifically, BIP-seq and TDOP-seq derived PPIs achieved accuracy rates of 62.5% (via BiFC) and 64.0% (via DLCA), respectively. Additionally, low-throughput pairwise validations yielded a 100% precision rate. For literature-derived data, POPPIN utilizes a DeepSeek-V4-Flash assisted text-mining workflow. Manual verification of a representative PPIs confirmed an 86.8% accuracy rate, further underscoring the platform’s high reliability (Supplemental Table S1). Finally, to evaluate the accuracy of POPPIN, we compared experiment-based rice PPIs in POPPIN with other reported databases, such as BioGRID, STRING, RiceNet, BioGRID, intact and MINT (Figure 1H; Supplemental Table 1).

## AUTHOR CONTRIBUTIONS

J. Z., JW. Z., and Z. E., designed and planned the research. X. L., J. L., L. J., D. X., and J. H., contributed equally to this work. X. L., J. L., L. J., D. X., J. H., Y. C., M. L., Y. C., X. L., G. L., W. L., J. L., J. Y., Y. W., Z. L., X. T., Y. H., performed experiments; X. L., J. L., and L. J., performed data collection. X. L., and L. J., trained the DeepSeek□V4□Flash models. X. L., J. L., and D. X., analyzed data and designed the database. X. L., J. L., L. J., J. Z., JW. Z., and Z. E., wrote and revised the manuscript. All authors read and approved the final manuscript.

## ACKNOWLEDGMENTS

The authors wish to express their gratitude to Prof. Qingyong Yang from Huazhong Agriculture University and Prof. Lida Zhang from Shanghai Jiao Tong University for their invaluable advice for the database design. The authors would also like to acknowledge the high-performance computing platform of National Key Laboratory of Crop Genetic Improvement at Huazhong Agriculture University and public lab of CNRRI.

---

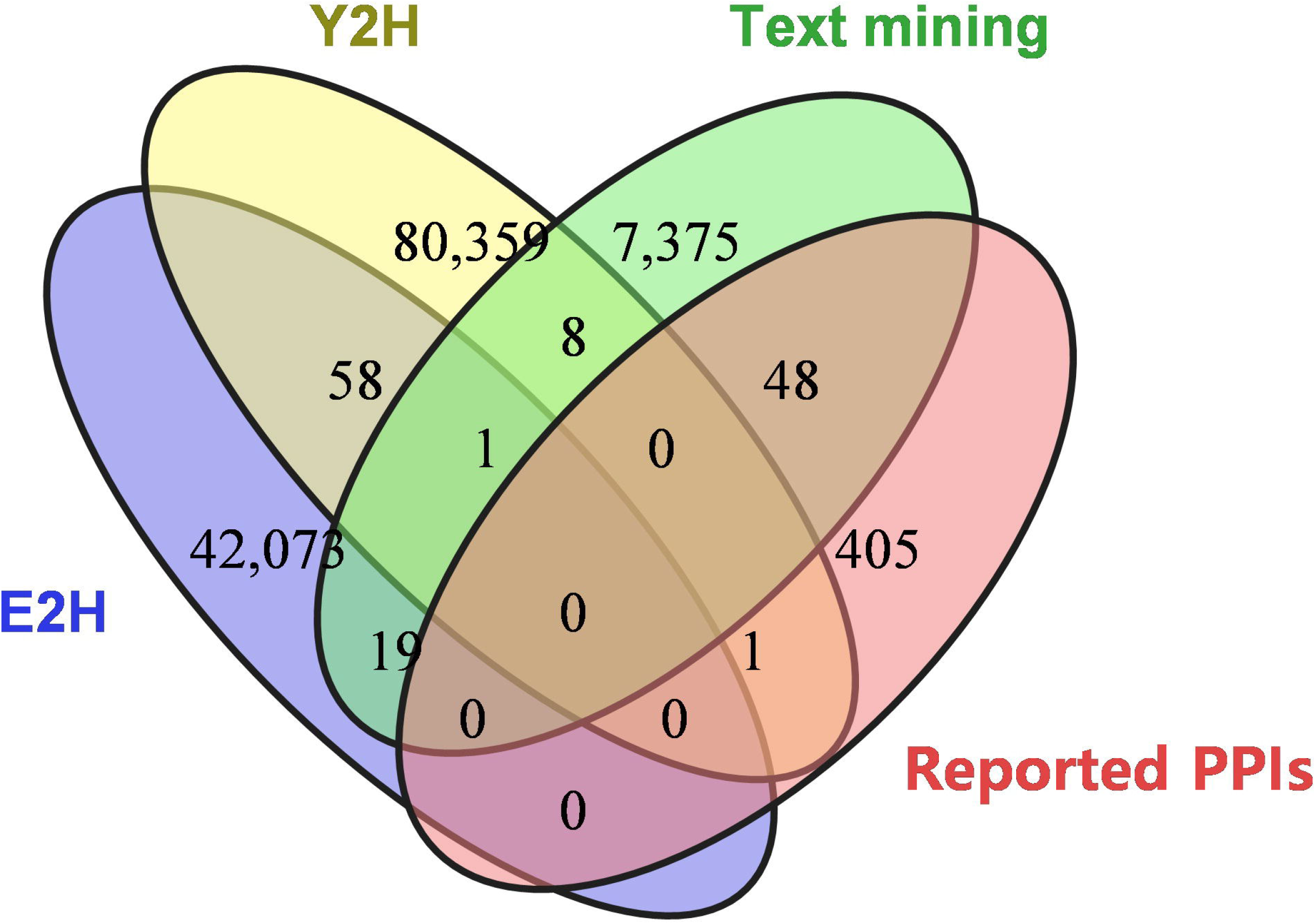

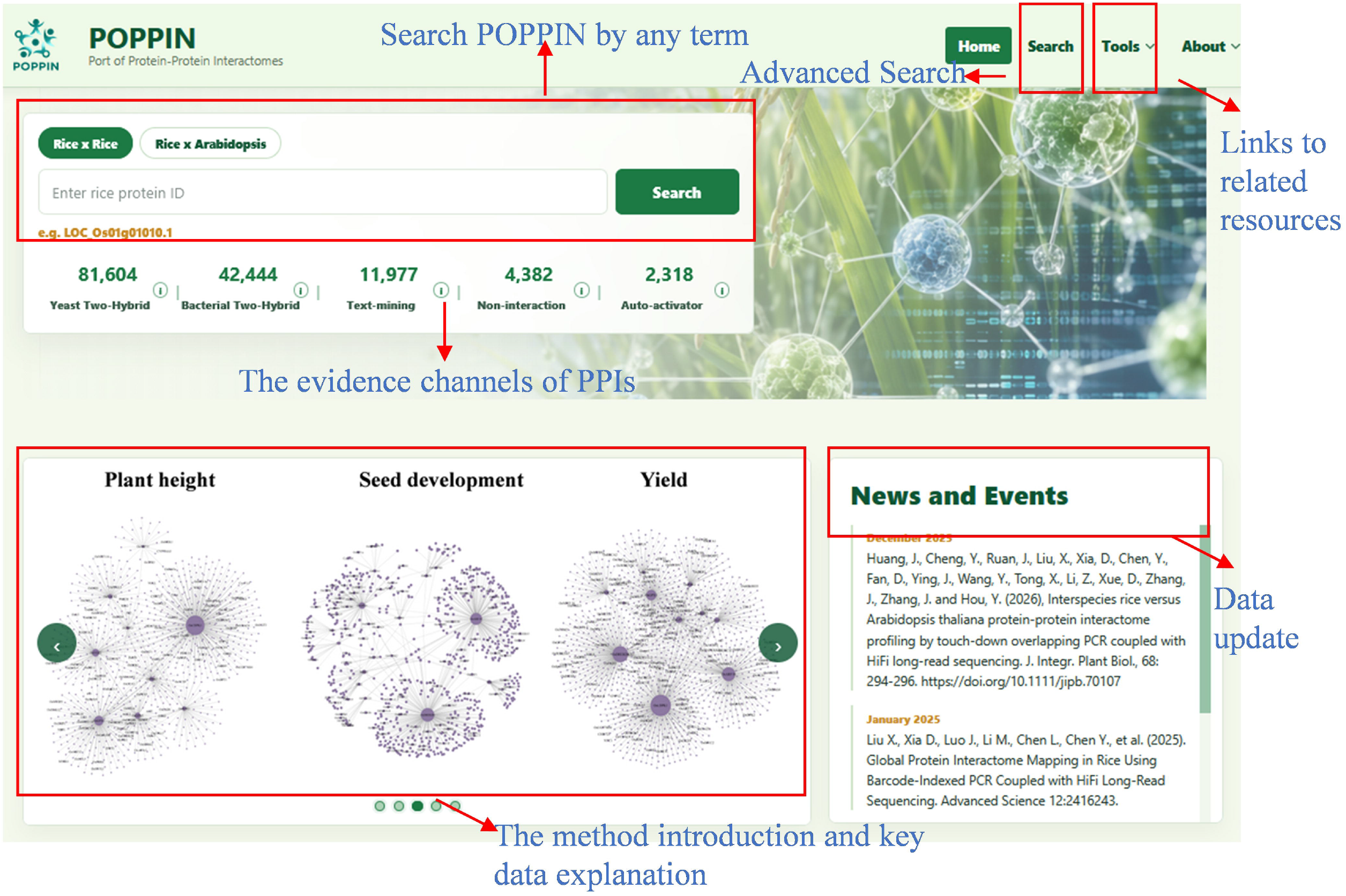

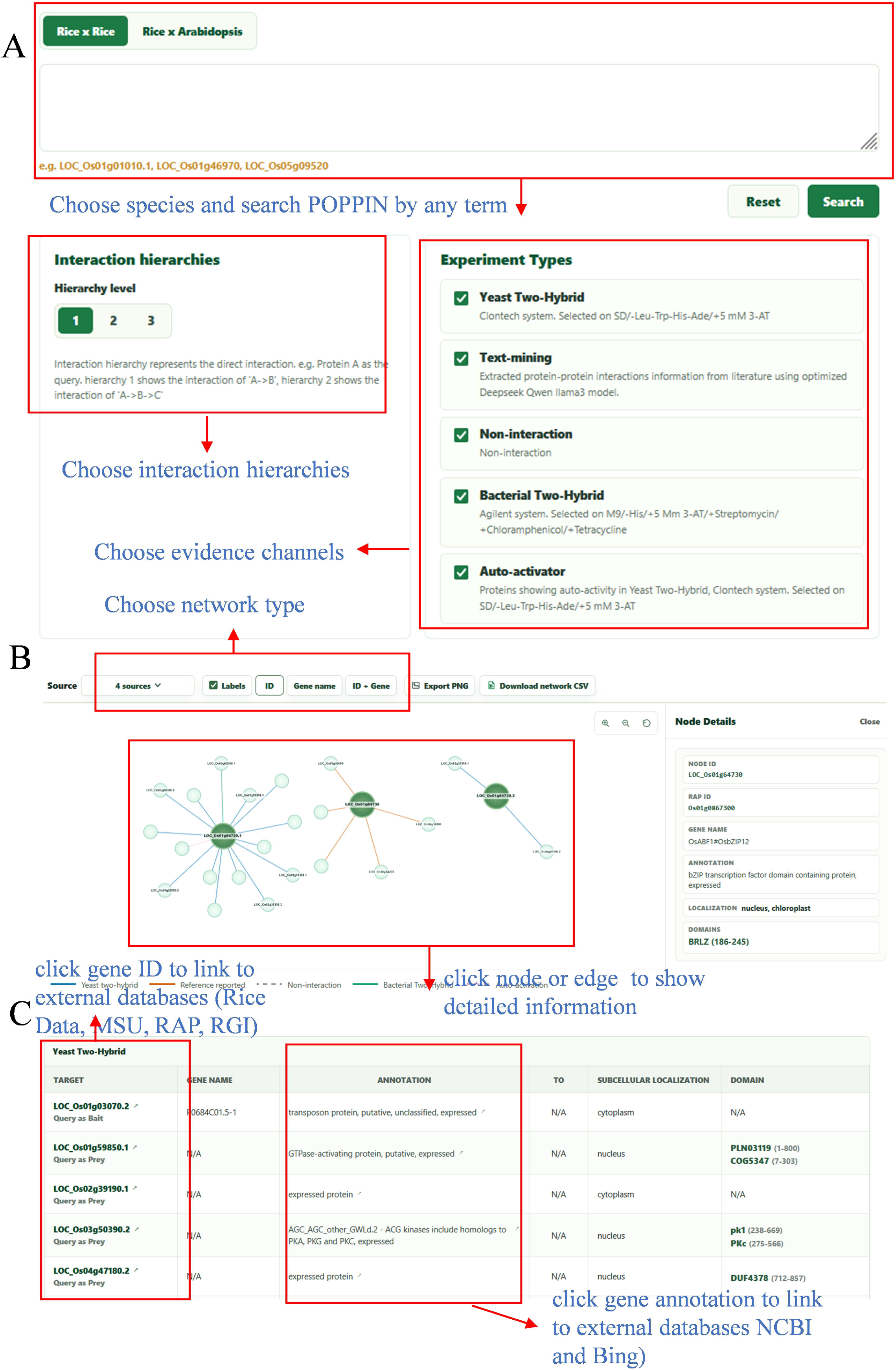

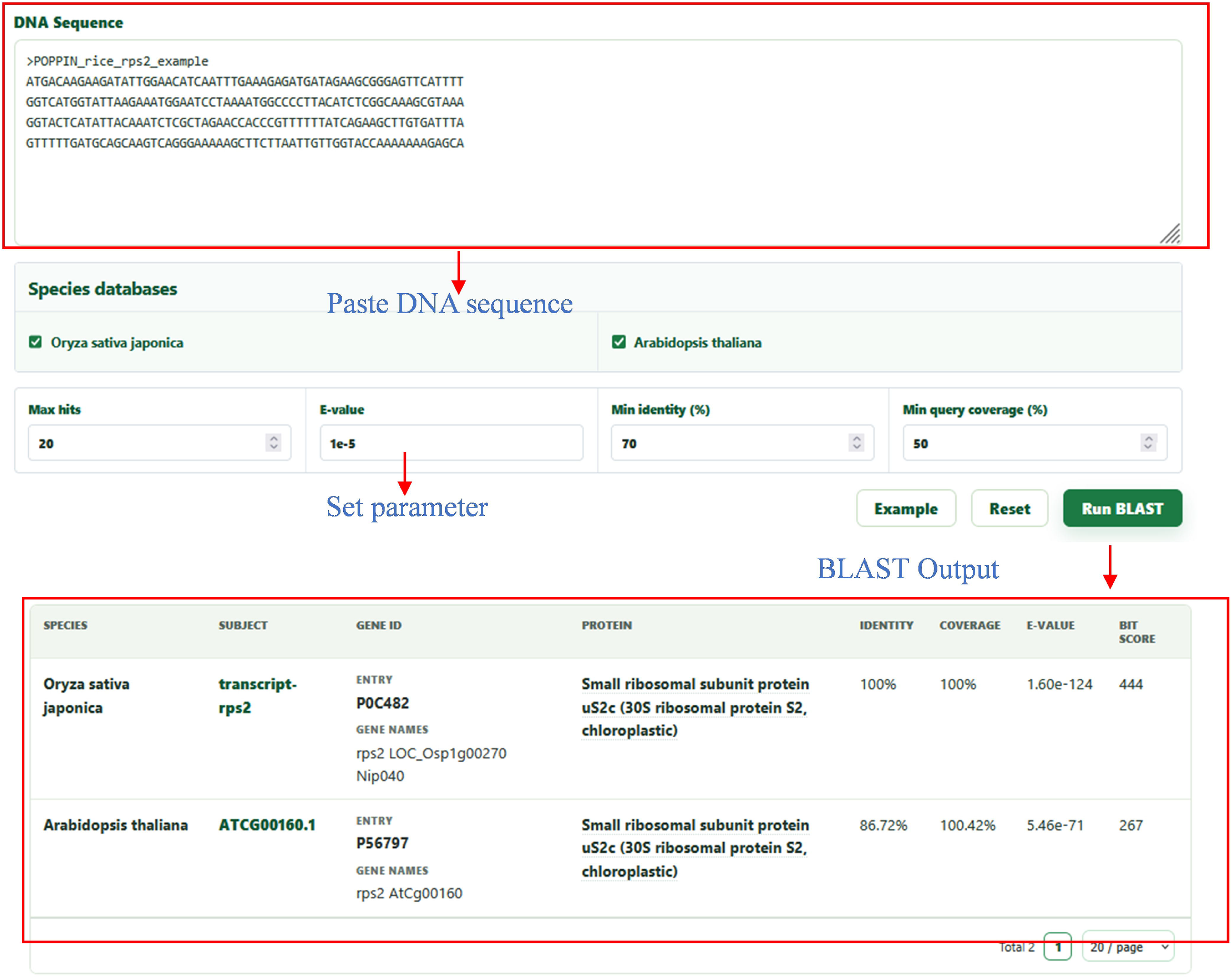

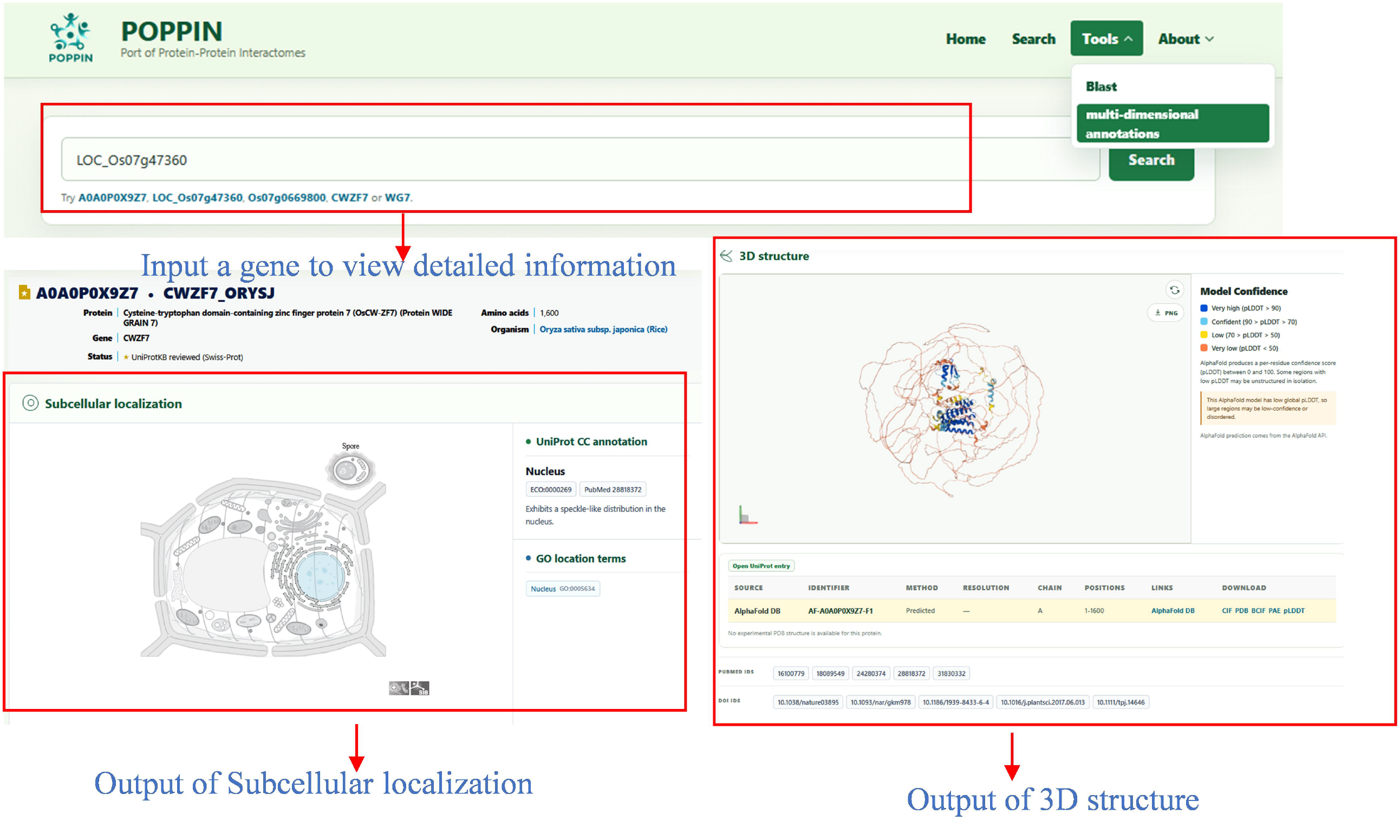

## Reference

Del Toro, N., Shrivastava, A., Ragueneau, E., Meldal, B., Combe, C., Barrera, E., Perfetto, L., How, K., Ratan, P., Shirodkar, G., et al. (2022). The IntAct database: efficient access to fine-grained molecular interaction data. Nucleic Acids Research 50: D648–D653. 10.1093/nar/gkab1006.

Deqing, L., Honghui, M., Yi, S., Shuang, S., Wenli, Z., Junting, W., Ming, Z., and Wei, C. (2018). ECharts: A declarative framework for rapid construction of web-based visualization. Visual Informatics 2:136–146. 10.1016/j.visinf.2018.04.11.

Fields, S., and Song, O. (1989). A novel genetic system to detect protein-protein interactions. Nature 340:245–246. 10.1038/340245a0.

Gu, H., Zhu, P., Jiao, Y., Meng, Y., and Chen, M. (2011). PRIN: a predicted rice interactome network. BMC bioinformatics 12:161. 10.1186/1471-2105-12-161.

E, Z.g., Yun, Z.J., Sheng, C.Y., Qian, Q., Lei, W., Hangzhou, China, Hangzhou, and China (2006). Information System of Rice Gene Database Based on INTERNET. Chinese Journal of Rice Science 29:653–657. 10.3969/j.issn.1001-7216.2015.06.012.

Hamilton, J.P., Li, C., and Buell, C.R. (2025). The rice genome annotation project: an updated database for mining the rice genome. Nucleic Acids Research 53: D1614–D1622. 10.1093/nar/gkae1061.

Huang, J., Cheng, Y., Ruan, J., Liu, X., Xia, D., Chen, Y., Fan, D., Ying, J., Wang, Y., Tong, X., et al. (2025). Interspecies rice versus Arabidopsis thaliana protein–protein interactome profiling by touch:LJdown overlapping PCR coupled with HiFi long:LJread sequencing. Journal of Integrative Plant Biology 68:294–296. 10.1111/jipb.70107.

Hui, Ge, and, Albertha, J.M Walhout, and, Marc, and Vidal (2003). Integrating ‘omic’ information: a bridge between genomics and systems biology - ScienceDirect. Trends in Genetics 19:551–560. 10.1016/j.tig.2003.08.009.

Kawahara, Y., Hirozane-Kishikawa, T., Hirata, R., Wang, X., Tamagaki, Y., Teramoto, Y., Tabei, N., Kumagai, M., Sakai, H., and Itoh, T. (2026). Rice Annotation Project Database (RAP-DB): Literature-Curated Gene Annotation and Integrated Omics Resources for Rice Functional Genomics and Molecular Breeding. Rice 19:51 10.1186/s12284-026-00924-6.

Lee, T., Oh, T., Yang, S., Shin, J., Hwang, S., Kim, C.Y., Kim, H., Shim, H., Shim, J.E., Ronald, P.C., et al. (2015). RiceNet v2: an improved network prioritization server for rice genes. Nucleic Acids Research 43: W122–127. 10.1093/nar/gkv253.

Liu, S., Liu, Y., Zhao, J., Cai, S., Qian, H., Zuo, K., Zhao, L., and Zhang, L. (2017). A computational interactome for prioritizing genes associated with complex agronomic traits in rice (Oryza sativa). The Plant journal 90:177–188. 10.1111/tpj.13475.

Liu, X., Xia, D., Luo, J., Li, M., Chen, L., Chen, Y., Huang, J., Li, Y., Xu, H., Yuan, Y., et al. (2025). Global Protein Interactome Mapping in Rice Using Barcode:LJIndexed PCR Coupled with HiFi Long:LJRead Sequencing. Advanced Science 12: e2416243. https://doi.org/1210.1002/advs.202416243.

Max, F., Lopes, C.T., Gerardo, H., Yue, D., Onur, S., and Bader, G.D. (2015) Cytoscape.js: a graph theory library for visualisation and analysis. Bioinformatics 32:309–311. 10.1093/bioinformatics/btv557

Oughtred, R., Rust, J., Chang, C., Breitkreutz, B.J., Stark, C., Willems, A., Boucher, L., Leung, G., Kolas, N., Zhang, F., et al. (2020). The BioGRID database: A comprehensive biomedical resource of curated protein, genetic, and chemical interactions. Protein Science 30:187–200. 10.1002/pro.3978.

Szklarczyk, D., Gable, A.L., Lyon, D., Junge, A., Wyder, S., Huerta-Cepas, J., Simonovic, M., Doncheva, N.T., Morris, J.H., Bork, P., et al. (2019). STRING v11: protein–protein association networks with increased coverage, supporting functional discovery in genome-wide experimental datasets. Nucleic Acids Research 47: D607–D613. 10.1093/nar/gky1131.

Yu, Z., Chen, Y., Zhou, Y., Zhang, Y., Li, M., Ouyang, Y., Chebotarov, D., Mauleon, R., Zhao, H., Xie, W., et al. (2023). Rice Gene Index: A comprehensive pan-genome database for comparative and functional genomics of Asian rice. Molecular Plant 16:798–801. 10.1016/j.molp.2023.03.012.

Zhang, Q., Li, J., Xue, Y., Han, B., and Deng, X.W. (2008). Rice 2020: a call for an international coordinated effort in rice functional genomics. Molecular Plant 1:715–719. 10.1093/mp/ssn043.

