## Supplementale information for "Port of Protein-Protein Interactomes: An experiment-based protein-protein interactome database for rice"

**Supplementary Figures**
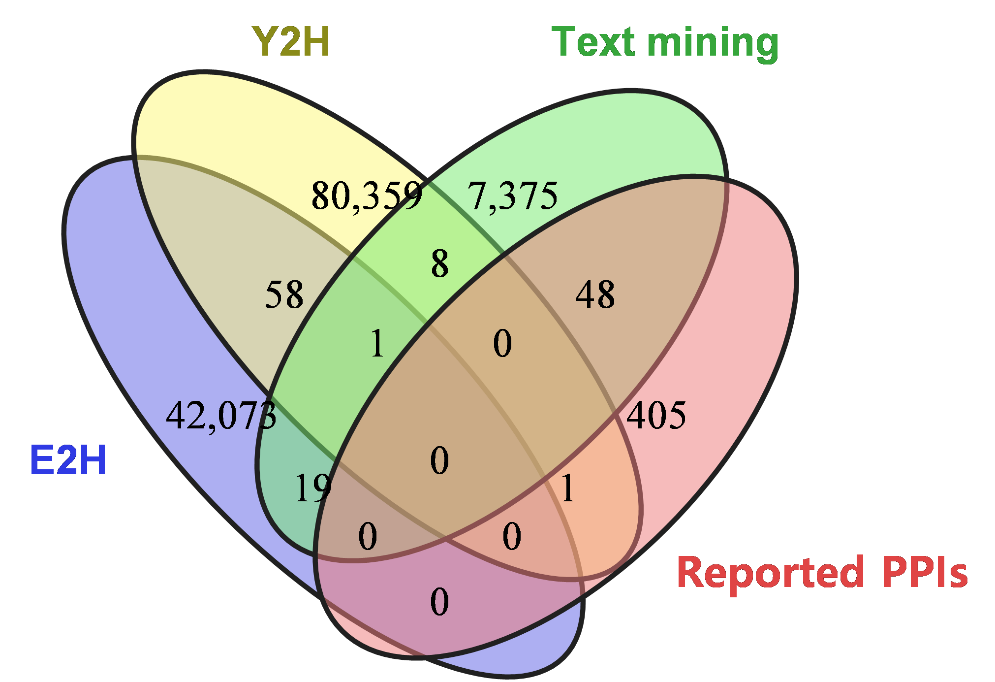

**Figure S1. Overlap between POPPIN with reported interaction datasets used for evaluating POPPIN quality**

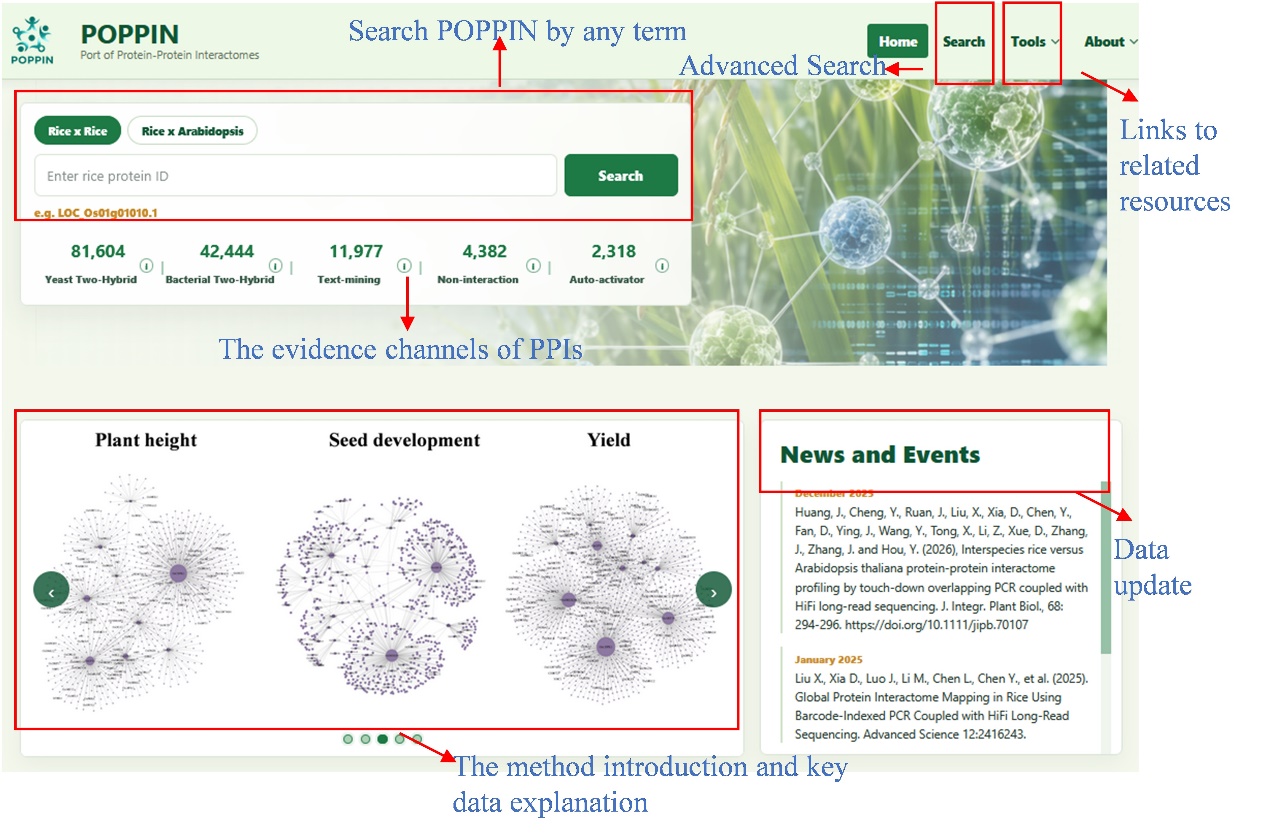

**Figure S2. Overview of the POPPIN database homepage**

The homepage displays a global search box, which users to quickly query intra-species (Rice × Rice) and cross-species (Rice × Arabidopsis) interactomes by any term. Below the search bar, quick-access sections display summarized statistics and provide direct navigation to major data categories. A dynamic scrolling banner shows methodological introductions and key data explanations (e.g., representative trait-specific network modules), alongside a "News and Events" panel tracking recent data updates and relevant publications.

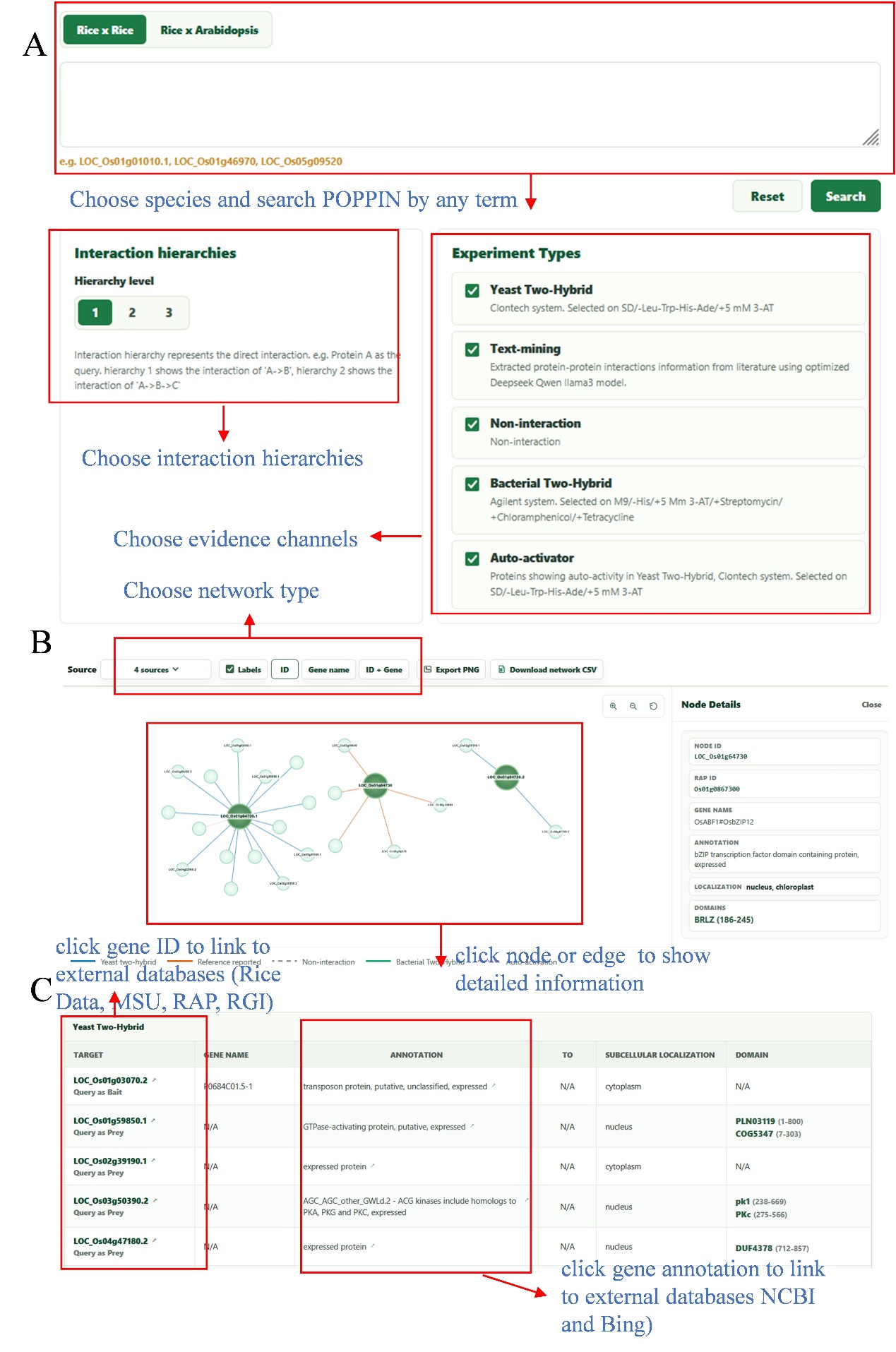

**Figure S3 Advanced search interface and interactive data visualization in POPPIN.**

(A) The advanced search interface. Users can perform multi-term queries across intra-species (Rice × Rice) or cross-species (Rice × Arabidopsis) interactomes. The interface allows filtering by defining "Interaction hierarchies" (hierarchy level from 1 to 3) and selecting specific "Experiment Types", including Y2H, B2H, Text-mining, and negative control datasets.

(B) Network visualization. PPI networks are rendered dynamically. Distinct edge colors and styles intuitively show different interaction evidence types (e.g., solid lines for Y2H/B2H, dashed lines for non-interactions). Clicking on specific nodes or edges displays the "Node Details" panel (right), displaying multi-dimensional annotations (e.g., RAP ID, subcellular localization, and conserved domains). The top toolbar enables users to customize node labels and export the network as a PNG image or CSV file.

(C) Table representation of interaction data. Search results are systematically organized in a detailed table format, summarizing target IDs, gene annotations, and corresponding functional properties. Both gene names and annotation provide the links with external databases such as MSU, Rice Data, RAP-DB, RGI, NCBI and Bing.

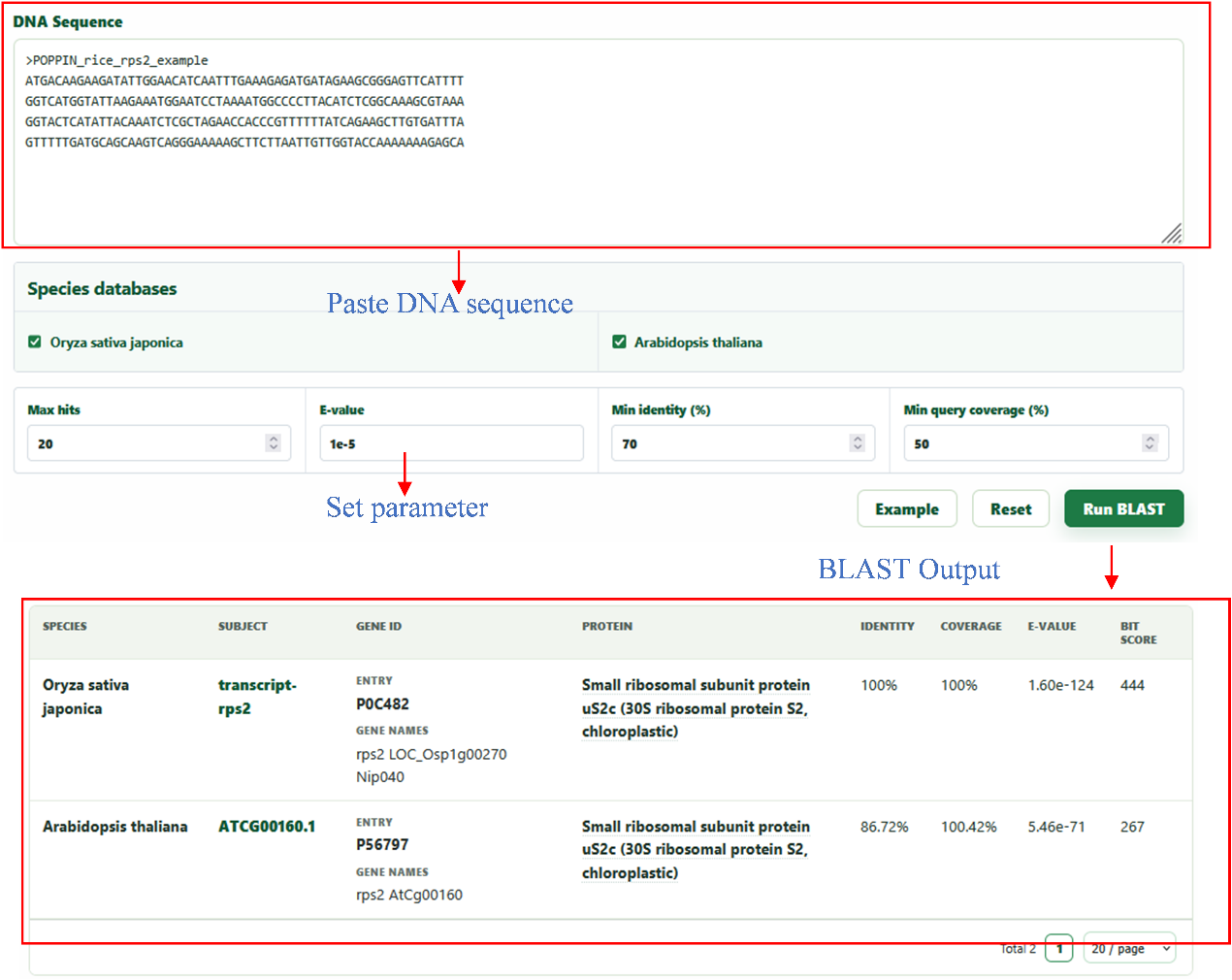

**Figure S4. POPPIN BLAST module: sequence similarity search**. The upper panel presents the BLAST query interface of the POPPIN database, designed to facilitate precise sequence similarity searches across its extensive gene collection. Users can seamlessly input nucleotide or protein uery sequences and select from specialized databases (e.g., cDNA, CDS, or protein sequences) while fine-tuning parameters such as E-value cutoff and alignment scores to optimize their search. Initiating the search via the "BLAST" button triggers a robust analysis. The lower panel exemplifies the resulting BLAST output report, delivering a table of sequence alignments. This includes a detailed table summarizing key metrics, such as query coverage, total score, E-value, and identity.

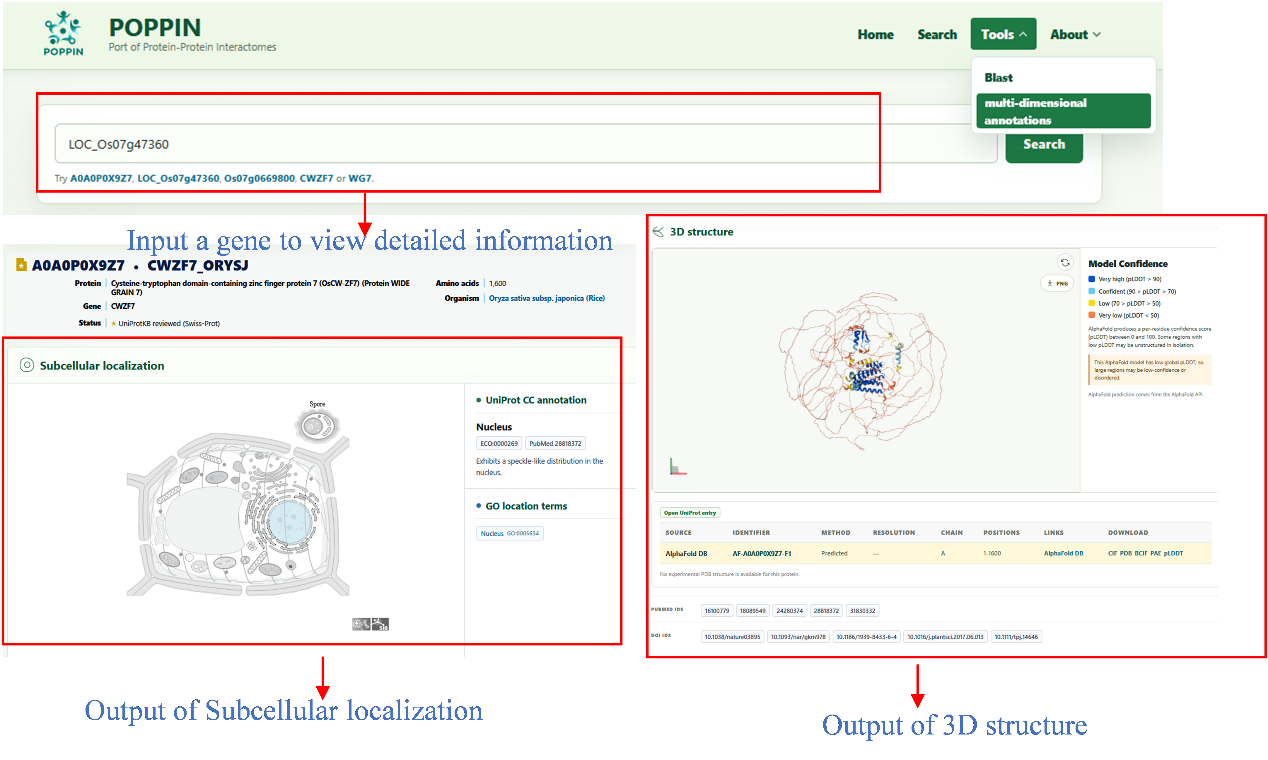

**Figure S5. POPPIN multi-dimentational annotation module: protein subcellular localization and 3D structure predition.** The upper panel presents the protein query interface of the POPPIN database, designed to predict gene Subcellular localization and 3D structure link to Uniprot dataset. Users can seamlessly input gene name or gene ID by initiating the prediction process with a single click of the “search" button. The lower panel exemplifies the resulting POPPIN output report, delivering comprehensive and precise functional predictions for each search protein. For each input protein, the results deliver a comprehensive visualization of subcellular localization and 3D structure. The result of subcellular localization confirms by uniport CC annotation and GO location terms on the right. Meanwhile, the color of 3D structure shows the model confidence. The results table presents details of source, identifier, method, resolution, chain, positions, links of this graphic.
